# A global to local genomics analysis of *Clostridioides difficile* ST1/RT027 identifies cryptic transmission events in a northern Arizona healthcare network

**DOI:** 10.1101/544890

**Authors:** Charles H.D. Williamson, Nathan E. Stone, Amalee E. Nunnally, Heidie M. Hornstra, David M. Wagner, Chandler C. Roe, Adam J. Vazquez, Nivedita Nandurkar, Jacob Vinocur, Joel Terriquez, John Gillece, Jason Travis, Darrin Lemmer, Paul Keim, Jason W. Sahl

## Abstract

*Clostridioides difficile* is a ubiquitous, diarrheagenic pathogen often associated with healthcare-acquired infections that can cause a range of symptoms from mild, self-limiting disease to toxic megacolon and death. Since the early 2000s, a large proportion of *C. difficile* cases have been attributed to the ribotype 027 (RT027) lineage, which is associated with sequence type 1 (ST1) in the *C. difficile* multilocus sequence typing (MLST) scheme. The spread of ST1 has been attributed, in part, to resistance to fluoroquinolones used to treat un-related infections, which creates conditions ideal for *C. difficile* colonization and proliferation. In this study, we analyzed 27 isolates from a healthcare network in northern Arizona, USA, and 1,352 public ST1 genomes to place locally-sampled isolates into a global context. Core genome, single nucleotide polymorphism (SNP) analysis demonstrated that at least 6 separate introductions of ST1 were observed in healthcare facilities in northern Arizona over an 18-month sampling period. A reconstruction of transmission networks identified potential nosocomial transmission of isolates following two of these introductions, which were only identified via whole genome sequence analysis. Antibiotic resistance heterogeneity was observed among ST1 genomes, including variability in resistance profiles among locally sampled ST1 isolates. To investigate why ST1 genomes are so common globally, we compared all high-quality *C. difficile* genomes and identified that ST1 genomes have gained and lost a number of genomic regions compared to all other *C. difficile* genomes; analyses of other toxigenic *C. difficile* sequence types demonstrates that this loss may be anomalous and could be related to niche specialization. These results suggest that a combination of antimicrobial resistance and gain and loss of specific genes may explain the prominent association of this sequence type with *C. difficile* infection cases worldwide. The degree of genetic variability in ST1 suggests that classifying all ST1 genomes into a quinolone-resistant hypervirulent clone category may not be appropriate. Whole genome sequencing of clinical *C. difficile* isolates provides a high-resolution surveillance strategy for monitoring persistence and transmission of *C. difficile* and for assessing the performance of infection prevention and control strategies.

## 1. Introduction

*Clostridioides difficile* (*1*) is one of the most commonly observed diarrheal pathogens in hospital settings. *C. difficile* infection (CDI) can range in severity from asymptomatic carriage or mild disease to toxic megacolon and death. The recent rise in the frequency in CDI has been attributed, in part, to the spread of fluoroquinolone-resistant strains of ribotype (RT) 027. Strain CD196, the earliest identified RT027 isolate (*2*), was isolated in France in 1985 (*3*). RT027 was linked to CDI outbreaks in North America and Europe in the 2000s (*4–8*) and has spread around the world (*9*). Although a decrease in the prevalence of RT027 has been reported in some regions (*10*), the RT027 lineage continues to be routinely isolated from clinical samples (*11–16*). Although the mechanisms behind the success of the RT027 lineage are not fully understood, the increase of CDI cases caused by RT027 in the 2000s has been linked to strains with fluoroquinolone resistance (*4, 9*). It has also been suggested that the RT027 lineage can displace endemic strains (*17*) and, more recently, Collins and colleagues (*18*) associated an increased ability to metabolize trehalose with epidemic ribotypes (RT027 as well as RT078) suggesting this trait along with the increased addition of trehalose to foods contributed to the success of RT027.

Many typing methods have been used to characterize *C. difficile* isolated from clinical settings. Ribotyping, a method that relies on differential amplification length profiles of ribosomal intergenic spacer regions, has been one of the most widely used procedures (*19, 20*). Multilocus sequence typing (MLST) has also been a commonly applied approach for characterizing *C. difficile* diversity and is more appropriate than ribotyping for examining evolutionary history and relatedness (*21*). RT027 is associated with sequence type (ST) 1 in the *C. difficile* MLST scheme (*22*). Additional ribotypes associated with ST1 include RT016, RT036, and RT176 (*21*). Ribotype 016 has been isolated from stool samples associated with CDI in England (*23*) and RT176 has been associated with CDI in Poland (*24*) and the Czech Republic (*25*). Ribotyping and MLST are increasingly being replaced by comparative genomic methods that rely on whole genome sequence (WGS) data. Ribotyping results, MLST profiles, and comparisons utilizing the entire genome are not always congruent (*20*); WGS provides the highest resolution for comparative genomics and should be the focus of comparative studies moving forward. The primary focus of this study is a comparison of ST1, which forms a monophyletic clade (see Results) and includes many virulent isolates collected from human clinical samples.

Multiple studies have investigated *C. difficile* with comparative genomics, with several studies focused on disease caused by ST1 (RT027). In one comparative study of three *C. difficile* genomes (*2*), the authors identified several genomic regions potentially associated with increased virulence of RT027. Since that time, genomic sequences from many additional strains have become available. A large comparative genomics study of *C. difficile* isolates collected from patients with diarrhea in hospital systems in the U.K. over a 4-year period indicated that the ST1 lineage was prevalent (17% of samples) (*26*). Another study that included 1,290 isolates from patients with CDI in the same area found that 35% of the isolates were ST1 (*27*). He and colleagues (*9*) investigated the global phylogeny and spread of RT027 (ST1) with whole genome SNP analysis and identified two epidemic lineages, both of which contained a mutation conferring fluoroquinolone resistance. As these studies demonstrate, ST1 isolates have been frequently identified from clinical samples; other studies have indicated that ST1 isolates are less commonly associated with other environments (e.g. soils, dogs, etc.) (*28–31*). Although not conclusive, together these studies suggest that ST1 isolates may preferentially colonize the human gut over other environments.

One of the primary concerns about CDI is emerging antimicrobial resistance (AMR). Antibiotic-associated pseudomembranous colitis has been associated with *C. difficile* since the late 1970s (*32–34*). Resistance to a variety of antimicrobials has been associated with *C. difficile* isolates, many of which are multi-drug resistant ((*35, 36*) and references therein). Antimicrobial resistance in *C. difficile* may vary between lineages or by region due to differing antimicrobial use and can impact CDI in regards to infection, recurrence, and disease outcome (*35, 36*). As mentioned previously, fluoroquinolone resistance has been associated with many ST1 (RT027) strains, and this resistance likely contributed, at least partially, to the increase in CDI cases attributed to ST1 during the 2000s as fluoroquinolone use increased during the 1990s and early 2000s (*4, 9, 37–39*). Fluoroquinolone resistance in ST1 strains has been associated with mutations in the *gyrA* and *gyrB* genes (*40, 41*). Suggested clinical guidelines for treating CDI currently include treatment with vancomycin, fidaxomicin, and metronidazole (*42*). Although reduced susceptibility to these compounds has been reported, resistance in *C. difficile* has had limited impact on the efficacy of these drugs in the clinic thus far (*42, 43*). Understanding AMR in *C. difficile* may yield insights into the prevalence of CDI caused by certain lineages and provide information regarding best practices for prescribing antibiotics.

In this study, we examined 27 ST1 genomes generated as part of a surveillance project across two healthcare facilities in northern Arizona, USA during 2016 and 2017 and publicly available *C. difficile* genomes. Comparisons were conducted to: 1) characterize northern Arizona ST1 isolates in the context of a worldwide set of genomes; 2) evaluate potential transmission networks within Arizona healthcare facilities; 3) use genomics characterize antimicrobial resistance profiles and to identify potential mechanisms that have allowed for the widespread presence and dominance of the ST1 lineage across hospitals worldwide; and 4) better understand the pangenomics and phylogenomics of the species in general. We also developed an *in silico* ribotyping method for relating whole genome sequence data and ribotyping information.

## 2. Methods

### Genome download and sequence typing

All available *C. difficile* genome assemblies (n=1,092) were downloaded from GenBank on November 30, 2017. For quality control purposes, statistics were gathered for each genome for number of contigs, number of ambiguous nucleotides (non A,T,G,C), and total genome assembly size. Additionally, all raw sequencing data (paired-end Illumina data only) associated with *C. difficile* available from the Sequence Read Archive (SRA) on September 1, 2017 were downloaded (sratoolkit fastq-dump). For isolates represented by multiple SRA runs, the runs were combined and labeled with the BioSample accession. Data were discarded if the read length was less than 75 bp. The multilocus sequence type (ST) of samples represented by raw read data was determined with stringMLST v0.5 (default settings) (*44*). Samples typed as ST1 were assembled with SPAdes v3.10.0 (--careful --cov-cutoff auto-k auto) (*45*). These genome assemblies were combined with GenBank assemblies for downstream analyses; in some cases, these reads represent duplicates of GenBank assemblies, but were treated independently in this study due to difficulties with association. Any GenBank assembly or assembly generated with SPAdes that contained greater than 1,500 contigs, greater than 10 ambiguous nucleotides, an anomalous genome assembly size (final data set: <3,600,277 or >4,698,454bp), or anomalous GC content (final data set: <28.07 or >29.74%) was removed from the data set. In addition, all pairwise MASH (v2.0, default sketch size) (*46*) distances were calculated for all genome assemblies in order to find genomes annotated as *C. difficile* but belonging to different species. Genomes with an average pairwise MASH distance >0.03, which corresponds to <0.97 average nucleotide identity (a conservative cutoff value), were removed from the dataset (average MASH distances for remaining genomes were less than 0.02). The ST was determined for all assemblies passing quality control metrics with a custom script (https://gist.github.com/jasonsahl/2eedc0ea93f90097890879e56b0c3fa3) that utilizes BLAST and the PubMLST database (https://pubmlst.org/) for *C. difficile* (*22*). For ST1 genomes, if the ST of the assembly did not match the ST predicted by stringMLST, the assembly was removed from the dataset. A total of 1,850 genome assemblies from NCBI data (609 Genbank assemblies and 1,241 in-house assemblies from the SRA identified as ST1) were included in this study (Table S1). Additionally, publicly available raw sequencing reads representing non-ST1 isolates (n=3,402) were included in the *in silico* ribotyping analyses (see below).

### *C. difficile* isolation, DNA extraction, sequencing, and assembly

Stool samples identified as containing *C. difficile* from two Northern Arizona Healthcare facilities (labeled facility A and facility B) were collected under IRB No. 764034-NAH to J. Terriquez and were stored at −80°C until processing. Isolation of *C. difficile* from stool samples was performed as outlined in Edwards *et al*. (*47*) with the following modifications; one 10 μL loopful of partially thawed stool was re-suspended in 500 μL of sterile 1x PBS in aerobic conditions. The suspension was immediately transferred to a vinyl Type C anaerobic chamber (Coy Laboratory Products, Grass Lakes, MI, USA), where 100 μL was plated onto pre-reduced taurocholate-cefotoxin-cyloserine-fructose agar (TCCFA) and incubated at 36°C for 24-48 hours. Suspected *C. difficile* were sub-cultured onto pre-reduced brain heart infusion agar supplemented with 0.03% L-cysteine (BHIS) and incubated anaerobically at 36°C for 24 hours. Once isolation of suspected *Clostridioides/Clostridium* spp. was achieved, a lawn was created (also on BHIS) and incubated anaerobically at 36°C for an additional 24 hours. Genomic DNA was extracted from each isolate, *C. difficile* species identification was confirmed via TaqMan^®^ PCR, and all *C. difficile* positive extractions were processed for downstream WGS as described previously (*31*). Based on this methodology, a total of 27 ST1 isolates were identified from hospitals in northern Arizona between March 2016 and September 2017 (Table 1). DNA was sequenced on the Illumina MiSeq platform and all genomes were assembled with SPAdes v3.10.0. The average per contig depth of coverage was calculated with genomeCoverageBed (*48*) from BWA-MEM v0.7.7 alignments (*49*). Additionally, 200 bases of each contig was aligned against the GenBank (*50*) nt database with BLASTN (*51*) and the identity of the top hit was tabulated. If the alignment was from a known contaminant or from a different species sequenced on the same run, the contig was manually removed from the assembly.

**Table 1.**
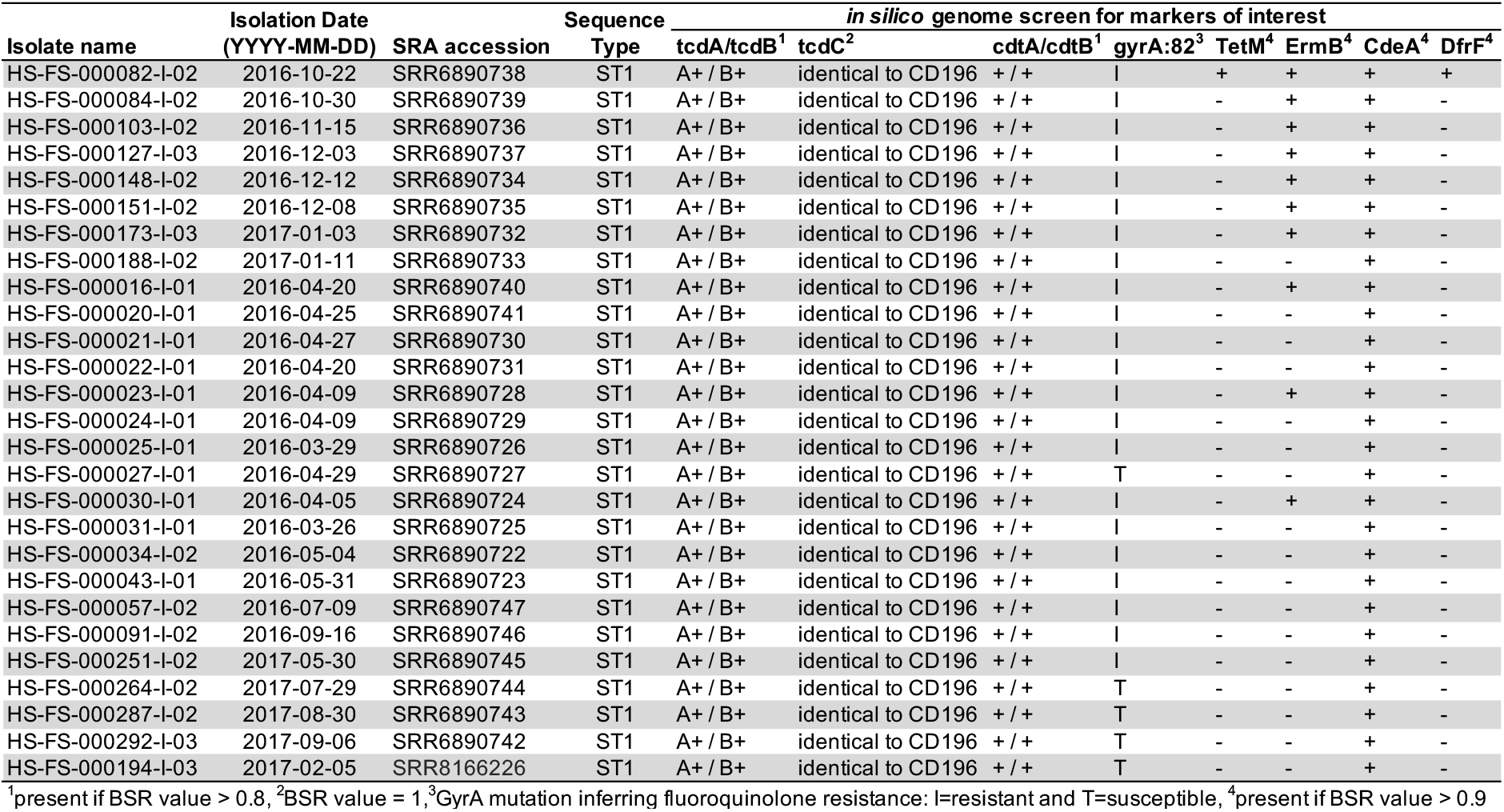
*C. difficile* ST1 isolates collected from a healthcare network in northern Arizona.

### Simulated reads

Genome assemblies have been shown to provide more noise when inferring the phylogenetic structure of a species compared to raw read data (*52*). For all publicly available genome assemblies downloaded from GenBank and passing filters, simulated reads were generated with ART (*53*) version MountRainier with the following command: art_illumina-ss MSv3 −l 250 -f 75 -m 300 -s 30. Simulated reads instead of genome assemblies were used for SNP discovery and phylogenetics for ST1 genomes.

### Phylogenetic analyses

To generate a preliminary phylogeny, genome assemblies (n=1,877) were aligned against the finished genome of *C. difficile* CD630 (GCA_000009205.1) with NUCmer (MUMmer v3.23) (*54*), and SNPs were called in conjunction with NASP v1.1.2 (*52*). A maximum likelihood phylogeny was inferred on an alignment of 85,331 concatenated SNPs (Table S2) called from a core genome alignment of 1,860,526 positions with IQ-TREE v1.6.1 (*55*) using the best-fit model (GTR+F+ASC+R5) identified by ModelFinder (*56*) and the UFBoot2 ultrafast bootstrapping option (*57*). Phangorn v.2.4.0 (*58*) was used to calculate the consistency index (excluding parsimony uninformative SNPs) and the retention index (using all SNPs) (*59*). All trees were visualized in FigTree v1.4.3 (http://tree.bio.ed.ac.uk/software/figtree/) or the Interactive T ree of Life online tool (*60*).

To generate a phylogeny for ST1 genomes (n=1,379), short read data (simulated reads for genome assemblies downloaded from NCBI and actual Illumina short reads for publicly available data and 27 newly sequenced genomes) were aligned against the finished ST1 genome CD196 (GCA_000085225.1) with BWA-MEM v0.7.7 (*49*). SNPs were called from the alignments with the Unified Genotyper method in GATK v3.3-0 (*61, 62*). Positions in duplicated regions of the reference genome (identified with NUCmer), positions with less than 10X coverage, and positions with a mixture of alleles (<0.9 single allele) were removed from the analyses. SNP positions with calls in at least 90% of the analyzed genomes were concatenated (n=4,283 SNPs, Table S3). A maximum likelihood tree was inferred with IQ-TREE v1.6.1 (best-fit model TVMe+ASC+R2). The tree was rooted with the ST1 genome ERR030325 based upon previous phylogenetic analysis. Phangorn was used to calculate the consistency index and retention index as described above.

### Root-to-tip regression and divergence time analysis

To estimate the date of divergence of the successful ST1 clade, a set of genomes (n=90; 87 ST1 genomes and 3 outgroup genomes) for which accurate sample dates were known were analyzed. SNPs were called with NASP (*52*) and the program PHIPack (*63*) was used to test for evidence of recombination, as this can confound divergence-dating analyses. To calculate accurate divergence times, recombination in ST1 SNPs was identified and removed using the program ClonalFrameML (*64*). Non-recombinatory SNP positions present in all ST1 genomes were processed further (Table S4). The presence of a temporal signal was assessed through regression analysis implementing root-to-tip genetic distance as a function of the sample year in the program TempEst version 1.5.1 (*65*) (http://tree.bio.ed.ac.uk/software/tempest/). The determination coefficient, R^2^, was used as a measure of clocklike behavior with the best-fitting root selected in an effort to maximize R^2^. Additionally, 10,000 random permutations of the sampling dates over the sequences were performed in an effort to evaluate the significance of the regression results (*66*).

The best nucleotide substitution model was inferred using the Bayesian information criterion in the software IQ-TREE version 1.5.5 (*55*). BEAST version 1.8.4 (*67*) was used to estimate evolutionary rates and time to most recent common ancestor (TMRCA) through a Bayesian molecular clock analysis using tip dating. BEAST analysis was run with a correction for invariant sites by specifying a Constant Patterns model in the BEAST xml file. The numbers constant As, Cs, Ts, and Gs were added to the BEAST xml file. A “path and stepping stone” sampling marginal-likelihood estimator was used in order to determine the best-fitting clock and demographic model combinations (*68*). The log marginal likelihood was used to assess the statistical fits of different clock and demographic model combinations (Table S5). Four independent chains of one billion iterations were run for the best clock and demographic model combination. Convergence among the four chains was confirmed in the program Tracer version 1.6.0 (http://tree.bio.ed.ac.uk/software/tracer/).

### *C. difficile* transmission and persistence analysis

To provide insight into persistence of *C. difficile* and transmission of *C. difficile* within and among healthcare facilities in northern Arizona, whole genome SNP phylogenies were coupled with epidemiological data. For northern Arizona isolate genomes within clades of interest, SNPs were called with NASP (reference CD196; GCA_000085225.1) and maximum parsimony phylogenies were inferred with Phangorn (*58*) (parsimony trees were used for easily mapping the number of SNP differences between isolates). Epidemiological data were collected from healthcare facilities but, in some cases, information was incomplete.

### *In silico* predicted antimicrobial resistance profiling

Proteins (n=2,177) from the comprehensive antimicrobial resistance database (CARD) (*74*) were downloaded on December 18th, 2017. Protein sequences were aligned against all ST1 genomes with the TBLASTN option in LS-BSR. Any peptide with a BSR (*75*) value > 0.9, which is equivalent to 90% ID over 100% of the peptide length (*76*), was investigated further.

Several mutations associated with fluoroquinolone resistance have been published in the literature (*36, 40, 41, 77*). For GyrA (CD630DERM_00060) and GyrB (CD630DERM_00050), predicted protein sequences were extracted for all genome assemblies from TBLASTN alignments and then aligned with MUSCLE (*73*). The states of mutations associated with fluoroquinolone resistance were manually investigated.

### Antibiotic resistance testing

Four ST1 isolates collected as part of this study (n=27) exhibiting various *in silico* predicted AMR profiles were screened for resistance to vancomycin, tetracycline, and ciprofloxacin on Brucella blood agar using Etests (bioMérieux, France). Inhibition ellipses were examined at 24 and 48 hours. Minimum inhibitory concentration breakpoints were based upon recommendations by CLSI (https://clsi.org/), the European Committee on Antimicrobial Resistance Testing (http://www.eucast.org/clinical_breakpoints/) and the available literature (*41*) and are as follows: vancomycin: <2 μg/ml - susceptible, 2-4 μg/ml - intermediate, >4 μg/ml - resistant; tetracycline: >16 μg/ml - resistant; ciprofloxacin: >16 μg/ml - resistant.

### Comparative genomics

For large-scale comparative genomics, genome assemblies were processed with the LS-BSR pipeline (*69*). In each genome coding regions (CDSs) were predicted with Prodigal v2.60 (*70*) and clustered with USEARCH v10.0.240_i86linux32 (*71*) at an identity of 0.9. A representative sequence for each cluster was then aligned against all analyzed genomes with BLAT v35x.1 (*72*). Scripts provided with the LS-BSR tool were used to identify core genome CDSs (pan_genome_stats.py) and compare BLAST score ratio (BSR) values of CDSs among groups of genomes (compare_BSR.py). To identify CDSs potentially differentially conserved among groups of interest, BSR values for individual CDSs were compared between genomes belonging to different STs (ST1: n=1379; ST8: n=31; ST15: n=34; and ST63: n=13). A CDS was considered conserved in a group (ST) if the BSR value for the CDS was greater than 0.8 for greater than 95% of target-group genomes and the BSR value was greater than 0.4 in less than 5% of genomes of the non-target group.

Gene regions have been identified in ST1 genomes that have been linked to virulence (*2*) (Table S6). Because these regions were discovered utilizing a small number of genomes, we screened the peptide sequences from these regions against all genomes with LS-BSR using the TBLASTN alignment option. Additionally, genome assemblies were screened for genomic features associated with trehalose metabolism (*18*) (Table S6). For TreR (CBA65726.1), sequences were pulled out of all genome assemblies from TBLASTN alignments and then aligned with MUSCLE (*73*). The state of a mutation associated with increased trehalose metabolism was manually investigated. Genome assemblies were also screened for proteins encoded by a four gene region associated with increased trehalose metabolism (FN665653.1:3231100-3237100) with LS-BSR using TBLASTN.

### *in silico* ribotyping

Standard ribotyping primers (5’-GTGCGGCTGGATCACCTCCT-3’, 5’-CCCTGCACCCTTAATAACTTGACC-3’) (*19*) were aligned against all completed ST1 genomes with an *in silico* polymerase chain reaction (PCR) script (https://github.com/TGenNorth/vipr) and the predicted amplicon products were identified. Seven amplicons of defined lengths (Table S7) were conserved across all completed ST1 genomes. To test the ability of these amplicons to differentiate genomes, raw sequencing reads representing 4,670 genomes were mapped against these 7 amplicons with Kallisto (*78*) using the “--bias” correction. For each sample, an amplicon was determined to be present if the read count for that amplicon was at least 20% of the maximum read count for any of the seven amplicons for that sample; this allows for a small amount of indiscriminate read counting.

For comparison to *in silico* results, PCR ribotyping was performed for 10 *C. difficile* samples representing multiple sequence types. Ribotyping PCRs were conducted using the same forward and reverse primers described in Janezic *et al*. 2011 (*79*). PCR reactions were carried out in 50 μL volumes containing the following reagents (given in final concentrations): 5-10 ng of gDNA template, 1x PCR buffer, 1.5 mM MgCl2, 0.2 mM dNTPs, 0.1 mg/mL BSA, 1.25 U Platinum^®^ *Taq* polymerase, and 1.0 μM of each primer. PCRs were cycled according to the following conditions: 95°C for 5 minutes to release the polymerase antibody, followed by 35 cycles of 95°C for 60 seconds, 57°C for 30 seconds, and 72°C for 60 seconds, followed by a final elongation step of 72°C for 10 minutes. We then conducted a PCR purification step aimed at removing primer dimer and unincorporated dNTPs using Agencourt AMPure XP beads (Beckman Coulter, Brea, CA, USA) at a 1:1 ratio (50 μL PCR product: 50 μL beads) according to the manufacturer’s recommendations with the following modifications: wash steps were conducted using 80% ethanol and final products were eluted in 25 μL of 0.01 M Tris-HCL buffer, supplemented with 0.05% Tween20. Electropherograms were generated for each of the PCR products using the Agilent 2100 Bioanalyzer (Agilent Technologies, Catalog # G2939BA), which provides greater resolution and specificity than a standard agarose gel. Specifically, the Agilent DNA 1000 kit (Agilent Biotechnologies, Catalog # 5067-1504) was used to analyze the sizing, quantity, and separation patterns of 1 μL for each of the DNA libraries.

### Genome accession information

ST1 genomes for isolates collected from healthcare facilities in northern Arizona were deposited in the National Center for Biotechnology Information Sequence Read Archive under BioProject accession # PRJNA438482 (https://www.ncbi.nlm.nih.gov/bioproject/PRJNA438482). Individual accession numbers are shown in Table 1.

## 3. Results

*C. difficile* ST1 isolates from two healthcare facilities in northern Arizona were examined to understand the ST1 isolates present in northern Arizona in the context of a global collection of *C. difficile*, to identify potential transmission events and to characterize AMR and the core/pan-genome within the isolates.

### Phylogenetic diversity of *C. difficile*

A maximum likelihood phylogeny (Figure 1) was inferred on a concatenation of 85,331 SNPs (Table S2) identified from a 1,860,526 nucleotide core genome alignment; the analysis included all *C. difficile* genome assemblies that passed through all quality filters (n=1,877; including 1,379 ST1 genomes). The consistency index (excluding parsimony uninformative SNPs) is 0.30, and the retention index is 0.97. It is important to note that not all currently described STs are represented by genome assemblies included in this analysis. For example, ST11 genome assemblies, which include RT078, were not included as they were filtered out based on large MASH distances. The results demonstrate that genomes identified as ST1, including all 27 isolates from northern Arizona, form a monophyletic clade. Other sequence types, such as ST2, ST3, and ST5 are paraphyletic, demonstrating the limitations of using sub-genomic sequence typing information to infer a common evolutionary history without consideration of WGS analysis.

**Figure 1.**
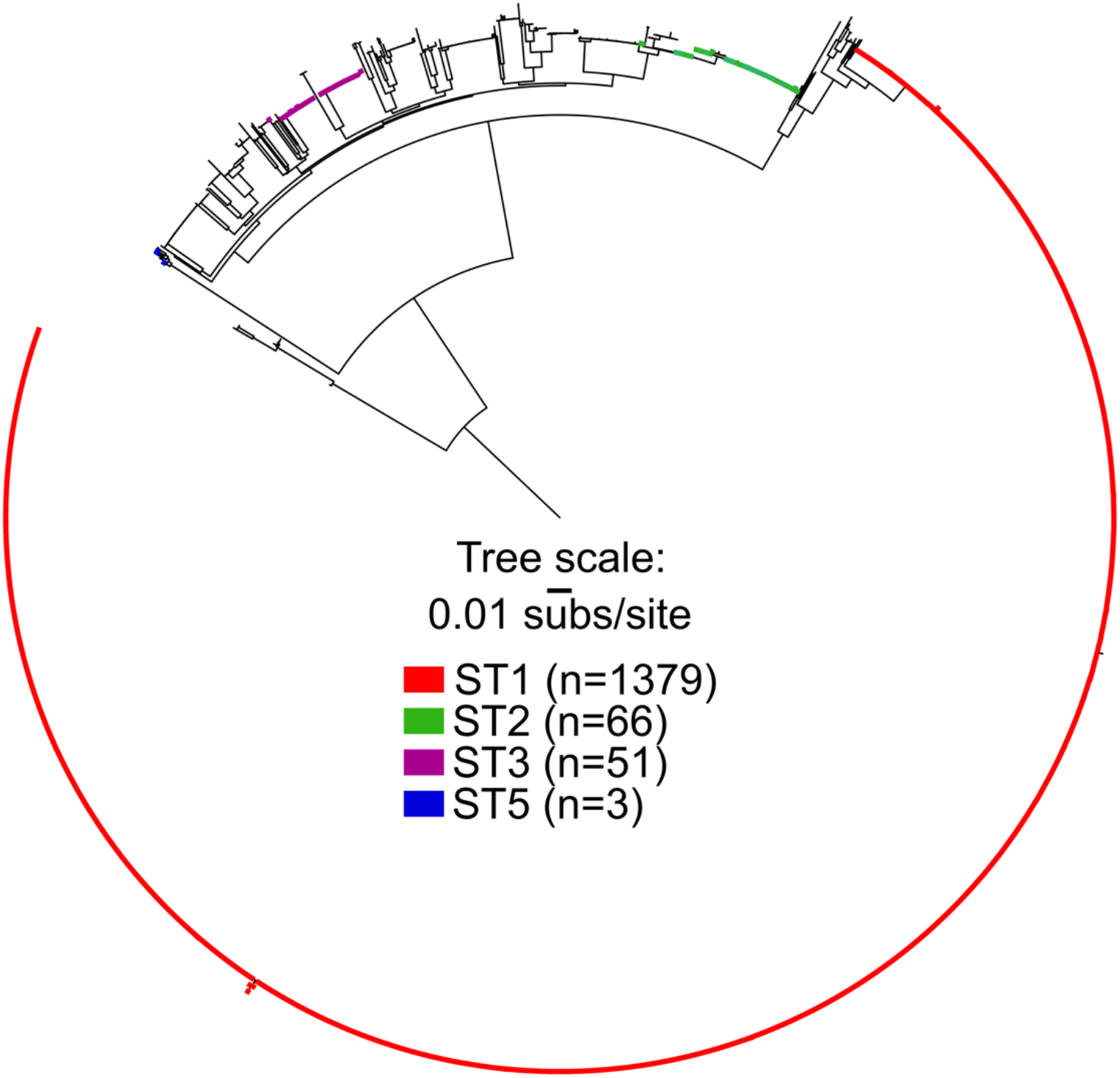
Maximum likelihood phylogeny of a global collection of *C. difficile* genomes (n=1,877) inferred from 85,331 SNPs showing the position of ST1 as well as other STs. ST1 forms a monophyletic clade (red) whereas other STs are paraphyletic (green, purple, blue).

An additional phylogenetic analysis was conducted for only ST1 genomes (n=1,379). A maximum likelihood phylogeny (Figure 2) was inferred on a concatenation of 4,283 SNPs (Table S3). The consistency index (excluding parsimony uninformative SNPs) is 0.87, and the retention index of this phylogeny is 0.98. The GyrA Thr82Ile mutation conferring fluoroquinolone resistance is conserved in two lineages labeled FQR1 and FQR2 (*9*). Isolates from two northern Arizona hospitals (n=27) that were sequenced as part of this study are found in FQR1 and FQR2 as well as lineages that do not have the GyrA Thr82Ile mutation. The northern Arizona isolates group into six independent clades (labeled 1-6 in Figure 2), separated by isolates collected from diverse geographic locations demonstrating that at least six separate introductions of ST1 isolates have occurred in the healthcare facilities over an 18-month time frame (Table 1, Figure 2). Clades that included more than one northern Arizona isolate (*1, 5* and *6*) were investigated further to determine the number of SNPs differentiating strains within each clade (pairwise comparisons of all high-quality positions in non-repetitive regions against CD196 reference). Northern Arizona isolates within clade 1, which also included isolates from other regions, vary by 0 to 31 SNPs (called from 3,617,365 genomic positions). Isolates from Arizona sequenced as part of previous studies (n=7) (*9, 80*) are present in two clades of the ST1 phylogeny, including clade 1. These previously sequenced isolates from Arizona are from human and food sources (*9, 81*), and pairwise comparisons indicate that the Arizonan isolates from previous studies within clade 1 of Figure 2 (collected from food samples in 2007) are separated from northern Arizona isolates (this study) by 8 to 22 SNPs (called from 3,617,365 genomic positions). Clade 5 includes two northern Arizona isolates collected from the same patient; the isolates had no SNP differences. Clade 6 is a monophyletic clade that includes only northern Arizona isolates; the genomes within this clade vary by 0 to 17 SNPs (called from 3,895,562 genomic positions). The average inter-clade (clades 1-6) SNP distance between northern Arizona isolates sampled in this study is 136 SNPs.

**Figure 2.**
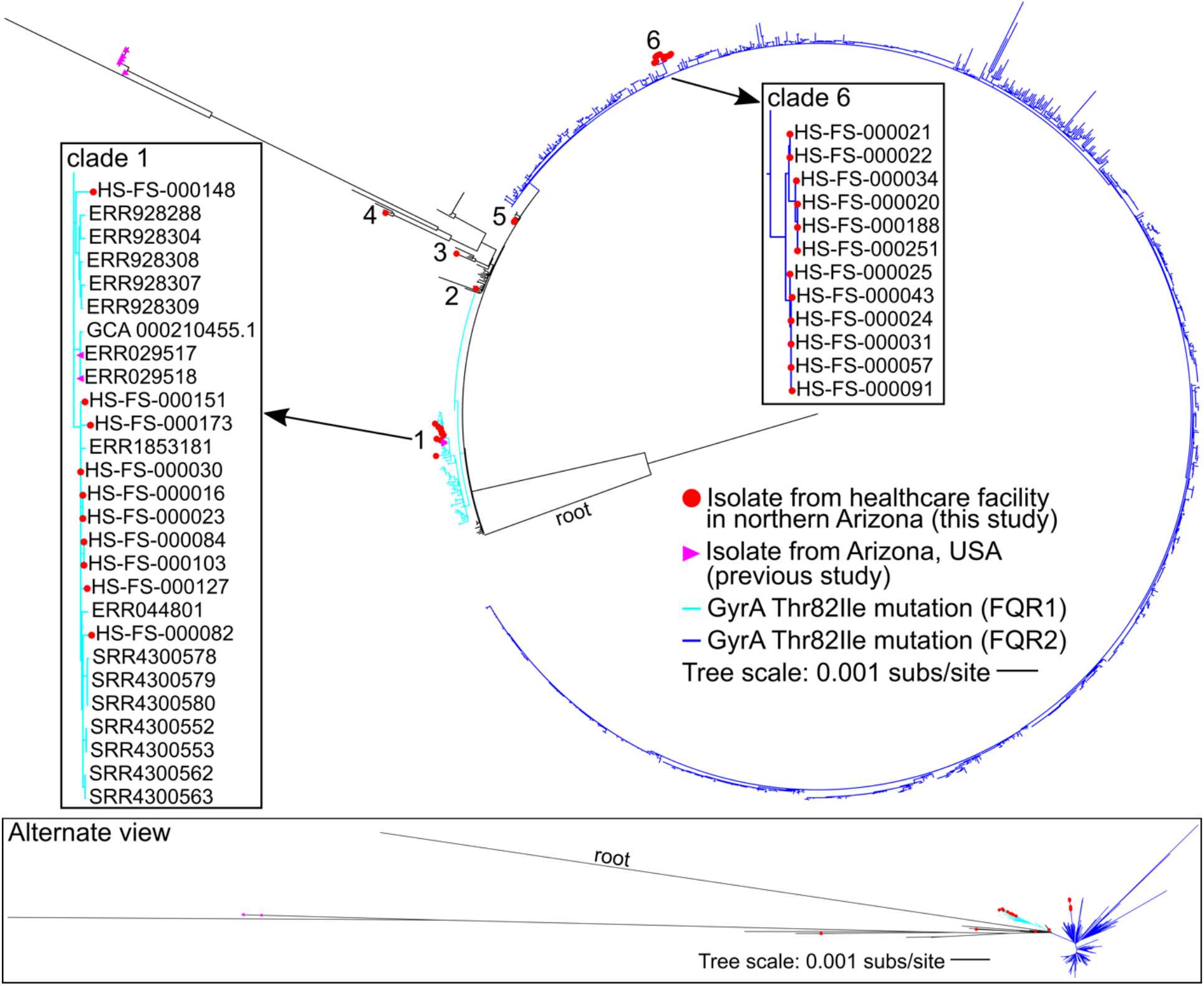
Maximum likelihood phylogeny of *C. difficile* ST1 genomes (n=1,379). Clinical isolates from healthcare facilities in northern Arizona (this study, in red) are present in six different clades (*1–6*). The presence of the GyrA Thr82Ile mutation conferring quinolone resistance is indicated by blue branches (light blue indicates FQR1 and dark blue indicates FQR2 identified in a previous study (*9*)). Seven isolates from Arizona sequenced as part of previous studies are identified with purple triangles. Boxes highlight clades (labeled 1 and 6) that contain multiple isolates from healthcare facilities in northern Arizona.

### Timing

Bayesian analysis of SNP data (with recombination removed) representing a set of ST1 and outgroup genomes (n=90) for which sample dates are known estimated the SNP accumulation rate at 1.13E-7 substitutions per site per year (highest posterior density (HPD) interval: 8.0632E-8 – 1.4634E-7). Considering the size of the *C. difficile* genome, this rate correlates to less than one SNP per genome per year. This SNP accumulation rate is slightly lower than a previous estimate for ST1 (*9*), as well as within host rates for *C. difficile* isolates from clinical samples (*26, 82*). The estimated divergence time for the ST1 clade genomes included in our analysis is 40.06 years ago (95% HPD interval between 32.14 to 49.54 years).

### Analysis of *C. difficile* transmission and persistence

Core genome SNP phylogenies for isolates collected from healthcare facilities in this study were paired with patient epidemiological data to provide insight into *C. difficile* prevalence, acquisition, and transmission within and between the two facilities (labeled A and B) (Figure 3). The estimated evolutionary rate within *C. difficile* ST1 genomes suggests that very few SNPs separate isolates involved in recent transmission events, which has been observed previously (*26*). Patient location information reveals that patients with CDI-associated diarrhea were frequently moved to multiple locations within a facility and once between facilities. Northern Arizona isolates within clade 1 are closely related, and isolates within clade 6 are also differentiated by very few SNPs (Figure 2). Isolates separated by 0 SNPs are observed at the same healthcare facility at multiple timepoints (e.g. Clade 6: HS-FS-000188, HS-FS-000020, HS-FS-000251 in Figure 3) and across both healthcare facilities (e.g. Clade 1 HS-FS-000016, HS-FS-000023; Clade 6: HS-FS-000024, HS-FS-000031, HS-FS-000057 in Figure 3). Isolates HS-FS-000057 and HS-FS-000043 are separated by 1 SNP and the two patients from which these isolates were sampled resided at the same skilled nursing facility, which is a third and separate entity from the two facilities at which samples were collected in this study. Thus, our analysis identifies potential transmission of *C. difficile* within the healthcare network. Some isolates (e.g. HS-FS-000148 in clade 1) are more distantly related to other sampled isolates, which could indicate infections acquired from community or environmental reservoirs. *C. difficile* within clades 1 and 6 appear to be prevalent in northern Arizona (though sampling in this study is limited to clinical specimens from two facilities) and are perhaps circulating within healthcare facilities.

**Figure 3.**
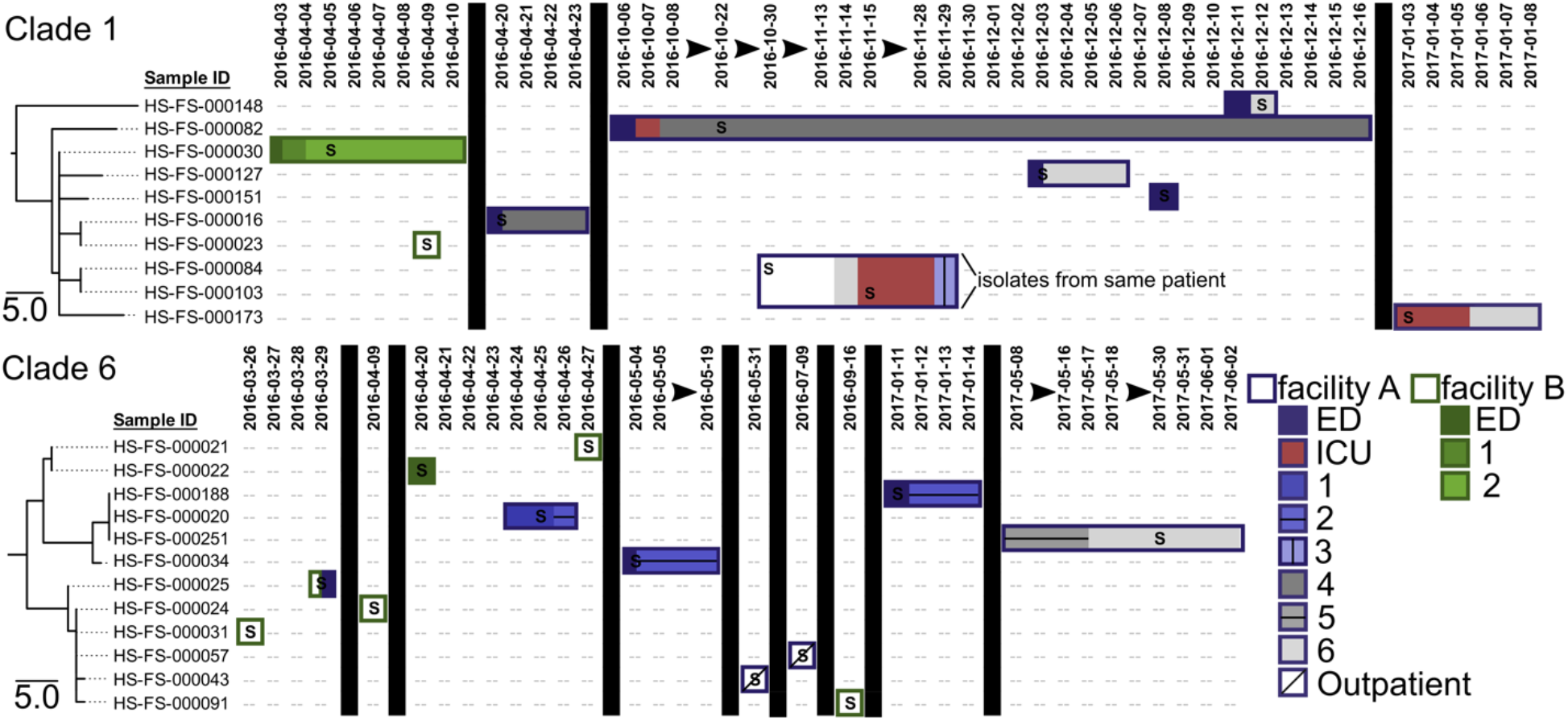
Maximum parsimony phylogenies for northern Arizona isolates within clades 1 and 6 identified in Figure 2 paired with patient location information to assess potential *C. difficile* ST1 transmission and persistence within and among facilities. *C. difficile* ST1 isolates originated from patient fecal samples collected at two facilities in the same healthcare network (facility A in purple, facility B in green). The location of the patient from which isolates originated is mapped over time (date formatting: YYYY-MM-DD) to the right of the phylogeny using rectangles. Patient locations at facility A are indicated with purple outlined rectangles whereas patient locations at facility B are indicated by rectangles outlined in green. Different wards within the facilities are indicated with differential shadings and patterns. Vertical black boxes and black arrows illustrate breaks in time. The letter “S” indicates the date of collection for the fecal sample from which each isolate originated. Isolates HS-FS-000084 and HS-FS-000103 originated from the same patient. The patient from which isolate HS-FS-000025 originated was transferred between facilities. The analysis identifies potential transmission of *C. difficile* within the healthcare network and persistence of a genotype within a patient.

On two occasions multiple isolates were sampled from the same patient. Isolates HS-FS-00084 and HS-FS-000103 (clade 1, Figure 2) were collected from the same patient 16 days apart; isolates HS-FS-000264 and HS-FS-000287 (clade 5, Figure 1) were collected from a different patient one month apart. In both cases, the isolates sampled from the same patient were separated by zero SNPs, indicating persistent infections or re-infections from the same source rather than new infections with a new genotype.

### *In silico* predicted AMR profiles in ST1 genomes

To understand AMR among northern Arizona isolates as well as ST1 in general, all ST1 genomes were screened against 2,177 proteins in the CARD database with LS-BSR (Table S8). The results indicate that although no proteins had hits (BSR value >0.9) across all ST1 genomes, several proteins were highly conserved across many ST1 isolates. The efflux transporter encoded by the *C. difficile cdeA* gene is highly conserved across ST1 genomes (average BSR value 0.99, BSR values > 0.9 in 1,375 of 1,379 ST1 genomes); this CDS is also present in all northern Arizona isolates (Table 1). Although over-expression of the protein in *E. coli* has been correlated with increased fluoroquinolone (norfloxacin and ciprofloxacin) resistance, the role of the CdeA protein in *C. difficile* is unclear (*91*). Some isolates such as CD196 have high BSR values for the CdeA protein but are susceptible to some fluoroquinolones (gatifloxacin, moxifloxacin, levofloxacin (*2*)) indicating the presence of this gene alone does not infer resistance to newer fluoroquinolones in ST1 genomes. Genes encoding proteins associated with TetM and ErmB genetic elements previously described to confer resistance to tetracycline and erythromycin in *C. difficile* (*36, 92–94*) are present in a number of ST1 genomes but are not universally conserved (TetM present in ~7% of ST1 genomes, ErmB present in ~8%). The *tetM* gene is present in one of the 27 northern Arizona isolates whereas the *ermB* gene is present in 10 northern Arizona isolates (Table 1). A gene encoding a dihydrofolate reductase (DfrF) protein was identified in 10 ST1 genomes including one isolate from northern Arizona (Table 1). DfrF has been associated with resistance to trimethoprim in *Enterococcus faecalis* (*95*) and *Streptococcus pyogenes* (*96*). Proteins encoded by the *vanG* operon associated with vancomycin resistance in *E. faecalis* (MIC 16 μg/ml) (*97*) were identified for two ST1 genomes (BSR values ranging from 0.92 to 1), including one isolate from northern Arizona (HS-FS-000151). These genes were identified in addition to the vanG-like gene cluster identified in *C. difficile* (*94*), which is highly conserved among *C. difficile* genomes.

In addition to gene presence/absence, several SNP mutations have been described that confer resistance to quinolones (*36, 40, 41, 77*). A comparison of SNP calls across all *C. difficile* genomes at those positions demonstrated that >95% of ST1 genomes contained the GyrA Thr82Ile mutation that confers quinolone resistance, whereas only ~12% of non-ST1 *C. difficile* genomes screened here have this mutation. Interestingly, genomic data for 5 of the 27 isolates from CDI cases at northern Arizona facilities indicate these isolates do not have the GyrA Thr82Ile mutation conferring quinolone resistance (clades 2-5 in Figure 2). Additional mutations in GyrA and GyrB associated with quinolone resistance were not broadly conserved in ST1 genomes (Table S9). These results suggest that AMR varies among ST1 isolates (and among northern Arizona isolates from the same hospital) and that factors in addition to AMR likely contribute to the success of ST1 in the human/hospital environment.

### Antimicrobial Resistance Testing

Four northern Arizona isolates (this study) with varied *in silico* predicted AMR profiles (Table 1) were screened for antimicrobial resistance using Etests (Table S10). These four isolates (HS-FS-000082, HS-FS-000127, HS-FS-000151, HS-FS-000264) were chosen to represent the diversity of *in silico* predicted AMR profiles among northern Arizona isolates (Table 1). As with most *C. difficile* isolates (*36*), all four isolates were resistant to ciprofloxacin (MIC >32 μg/ml), which is a commonly prescribed fluoroquinolone. One isolate (HS-FS-000082) was also resistant to tetracycline (MIC >32 μg/ml) and this isolate contained the *tetM* gene (see Results section *In silico* predicted AMR profiles in ST1 genomes). Three of the four isolates had intermediate resistance (MIC 4 μg/ml) to vancomycin as has been observed in other *C. difficile* isolates (*98, 99*). One of these isolates (HS-FS-000151) with intermediate vancomycin resistance contained CDSs associated with a *vanG* operon in *E. faecalis* (described above); however, this region did not appear to confer increased resistance to vancomycin in the tested isolate.

### Pan-genome composition of ST1 genomes

LS-BSR analyses were performed to understand the size and extent of the ST1 core and pan-genome, to identify CDSs that may contribute to the association of ST1 with the hospital environment, and to evaluate potentially unique features of northern Arizona isolates. The strict core genome (CDSs conserved in all genomes using the LS-BSR method) of 1,379 *C. difficile* ST1 genome assemblies was determined by LS-BSR to be 1,991 CDSs (Table S11), which corresponds to approximately 55% of the total CDSs in the ST1 genome CD196. The analyzed ST1 genomes spanned a wide range of assembly quality, which could result in underestimation of the core genome size. However, evaluating CDSs that are highly conserved among ST1 genomes could provide insight into why the lineage is so successful in addition to fluoroquinolone resistance. BSR values were used to identify CDSs that have been gained or lost in ST1 genomes. CDSs (n=44) were identified as highly conserved among ST1 genomes and largely absent from non-ST1 genomes, whereas 14 CDSs were identified as being lost from ST1 genomes (highly conserved among non-ST1 genomes and largely absent from ST1 genomes, see methods for criteria) (Table S12). CDSs identified as highly conserved in ST1 include some genomic regions previously identified as unique to RT027 isolates in a comparison of three genomes (*2*). For example, CDSs associated with the insertion of a catalytically more efficient *thyA* gene that disrupts the *thyX* gene in previously studied RT027 isolates (*83*) are highly conserved in the ST1 genomes analyzed here; these CDSs were upregulated by an ST1/RT027 isolate during the early stage of an infection in a monoxenic mouse model (*84*). Additional CDSs highly conserved in ST1 genomes are annotated as transposases or have unknown functions (Table S12). Binary toxin genes (*cdtA* and *cdtB*) are highly conserved within ST1 and largely absent from many other STs but did not meet the threshold defined here to be included as gained/lost CDSs.

To understand if the phenomenon of gene acquisition/loss is common across all clades of *C. difficile*, a comparable analysis was conducted on ST8 (RT002 and RT159) genomes (n=31). The results demonstrate that although no CDSs are unique to ST8, 30 CDSs are differentially conserved in ST8 and two CDSs are generally deleted (see methods for criteria). Genomes (n=13) from ST63 (RT053), which have been shown to contain similar *in silico* predicted AMR profiles to ST1 (see below), were also compared to understand CDS conservation and loss. The results demonstrate that 26 CDSs are highly conserved among ST63 genomes and no regions appear to be specifically lost by this lineage. For genomes (n=34) identified as ST15 (RT010), which includes non-toxigenic strains, 118 CDSs were identified as highly conserved and 26 CDSs were identified as generally deleted. These analyses may suggest that ST15 and ST1 are adapting to different niches/environments. ST1 is likely adapting to the human gut, and a large percentage of isolates from patients with CDI in northern Arizona belong to ST1.

Although ST11 (RT078) genomes were excluded from the initial round of this analysis due to filtering genome assemblies by MASH distance, CDSs identified as highly conserved or generally lost within ST1 were compared to ST11 lineage genomes (n=15). The ST11 lineage has been associated with severe disease and has been commonly isolated from clinical samples as well as agricultural, animal, and retail meat samples (*81, 85–90*). Several CDSs identified as highly conserved within ST1 were also conserved in ST11 genomes (Table S12). These CDSs include a DNA-binding regulator, a histidine kinase, and an RNA polymerase sigma factor. Some CDSs identified as generally lost within ST1 are conserved within ST11 (e.g. a TetR/AcrR family transcriptional regulator).

Although LS-BSR analysis indicates that over half of the predicted CDSs present in the genome for strain CD196 are part of the strict core genome for ST1 isolates, some of the CDSs present in this genome (and others within ST1) are differentially conserved throughout the ST1 lineage. Some CDSs are clade-specific or have been lost in particular clades of ST1 (Figure S1). 5,937 CDSs were identified in the accessory genome for ST1, and 1,049 CDSs were identified as unique to one genome. This variation in genomic content within the ST1 lineage suggests that different clades/isolates within ST1 will vary phenotypically, which could impact AMR, transmission, and virulence. For example, Stone and colleagues (submitted and in review) found variable toxin production among three ST1 strains using a transepithelial electrical resistance (TEER) assay. To understand the genetic composition of northern Arizona isolates, LS-BSR was used to identify any unique CDSs within clades 1-6 in Figure 2 compared to other ST1 isolates, but no CDSs were found to be unique to any of these clades. However, isolates HS-FS-000082 and HS-FS-000151, both within clade 1, each contained a CDS not identified in any other ST1 genome (HS-FS-000082 – ABC transporter permease, HS-FS-000151 – site-specific integrase).

A set of 52 peptide sequences (*2*) previously associated with differential conservation in ST1 genomes (Table S6) were screened against all genomes with LS-BSR. The results indicate that several features associated with an epidemic ST1 strain are also present in non-ST1 genomes (Table S13) or are not highly conserved among other ST1 genomes. Although this result does not rule out that these regions are associated with virulence of some ST1 strains, it does suggest that these regions do not fully explain the success of the ST1 lineage. Collins and colleagues (*18*) identified genomic features associated with increased metabolism of trehalose in epidemic ribotypes. A mutation in the TreR protein (Leu172Ile) associated with trehalose metabolism is present in predicted proteins for 1,375 of 1,379 screened ST1 genome assemblies. Truncated predicted proteins were identified for two ST1 genome assemblies (ERR030337, GCA_900011385.1), and predicted proteins with low identity to the TreR protein for *C. difficile* strain CD196 (CBA65726.1) were identified for two genome assemblies isolated from northern Arizona (HS-FS-000084, HS-FS-000103; clade 1 in Figure 2); these two isolates have a smaller genome (~100Kb) than other genomes and originated from the same patient over 16 days. Four protein sequences associated with trehalose metabolism in RT078 (ST11) isolates (*18*) were also screened against genome assemblies. High BSR values (>=0.96) for all four protein sequences were identified for genome assemblies representing multiple MLST sequence types (Table S13), including three ST1 genome assemblies (ERR026353, ERR030384, ERR251831; in-house assemblies from publicly available sequencing read data). The four protein sequences were not identified for any of the 27 northern Arizona isolates sequenced here.

### *In silico* ribotyping

The gold standard for *C. difficile* genotyping has been ribotyping, wherein RT027 isolates are primarily associated with ST1 (*22*). Here we have focused on ST1, which forms a monophyletic clade within a whole genome SNP phylogeny (Figure 1). However, there appears to be a disconnect in the literature between researchers using ribotyping and those using either MLST or WGS approaches. To relate northern Arizona isolates to published *C. difficile* PCR ribotypes without the need for performing PCR ribotyping in the laboratory, we explored the potential to extract ribotyping profiles from sequence data. To examine the utility of an *in silico* ribotyping approach, raw sequencing reads from the SRA with associated MLST profiles were aligned against amplicons predicted by probing standard ribotyping primers against three finished ST1 genomes, and hits were identified from samples with even read mapping across all amplicons (see methods). An analysis of different thresholds for variable read mapping demonstrated that 20% of the maximum read counts was an appropriate threshold for calling ribotype amplicons (Table S14). Of the 4,670 genomes screened, 1,268 of them were sequence typed as ST1 using stringMLST, and 1,253 of these ST1 genomes were identified as RT027 based on *in silico* ribotype profiles (Table S15). The *in silico* ribotyping method correctly identified ~99% of ST1 genomes based on their ribotype profiles. Only 17 of 3,402 non-ST1 genomes were identified as ST1. It is assumed here that all ST1 genomes analyzed are RT027 and all non-ST1 genomes are not RT027. ST1 also includes RT016, 036 and 176, but predominantly RT027 isolates have been associated with ST1. All 27 northern Arizona isolates were *in silico* ribotyped as RT027 and sequence typed as ST1 (*in silico*) according to the MLST scheme for *C. difficile*.

PCR ribotyping was performed on 10 *C. difficile* isolates with various sequence types for comparison to *in silico* results (Figure S2). Two ST1 isolates presented similar PCR ribotyping profiles, and PCR ribotyping profiles for these two ST1 isolates were comparable to *in silico* ribotyping results, though not exact matches. The *in silico* profile included seven bands; one ST1 isolate presented analogs to all seven bands plus one larger band whereas the other ST1 isolate presented six analogous bands plus one larger band (Table S16). Variation between ribotyping profiles predicted from four RT027 genome assemblies and actual ribotyping profiles has been reported previously (*100*). Non-ST1 isolates produced dissimilar PCR ribotyping profiles to ST1 isolates, and very few non-ST1 isolates were identified as ST1 by the *in silico* ribotyping method. The *in silico* ribotyping method described here could potentially provide a means to connect ribotyping information from past studies to the expanding whole genome sequence data available for *C. difficile* isolates, though the method requires further testing and validation.

## 4. Discussion

*C. difficile* ST1 is a successful worldwide lineage that appears to be especially adapted to the human gut and healthcare environments. The rise of *C. difficile* in hospitals globally has been attributed, at least partially, to the emergence and proliferation of ST1 (RT027) (*101*). In this study, we sequenced 27 ST1 isolates collected between March 2016 and September 2017 from two healthcare facilities in northern Arizona (these 27 isolates represent ~15% of all isolates collected as part of a larger surveillance study) and compared them to a worldwide collection of *C. difficile* ST1 genomes. The results demonstrate that diverse ST1 isolates were present in the sampled healthcare facilities; northern Arizona isolates are distributed throughout the ST1 phylogeny, including within two previously identified fluoroquinolone resistant lineages (FQR1 and FQR2 (*9*)). At least six separate introductions (clades) of *C. difficile* ST1 into the two sampled healthcare facilities were observed over the period of surveillance (Figure 2). Four of the introductions/clades are represented by only one isolate or genotype (clades 2-5 in Figure 2), whereas clades 1 and 6 contain multiple northern Arizona genotypes (defined as any SNP variation among isolates). Clade 6 is a distinct lineage that includes only isolates from northern Arizona facilities. Interestingly, two isolates collected from food samples in Arizona as part of previous studies (*9, 81*) are closely related to some of the clinical isolates sequenced in this study (clade 1 in Figure 2). *C. difficile* within clades 1 and 6 seem to be prevalent ST1 genotypes circulating within northern Arizona.

Whole genome sequencing provides a high-resolution method for differentiating bacterial isolates and offers insight into how pathogens persist and are transmitted within healthcare networks. Several studies, including this one, have indicated that the SNP accumulation rate in *C. difficile* is approximately one SNP per genome per year (*26, 82*); however, SNP accumulation rates can be complicated for bacteria with a spore stage in their life cycle (*102*). Bayesian analysis suggests that the divergence time for the ST1 lineage is approximately 40 years ago (95% HPD interval between 32.09 to 49.46 years), which is consistent with the identification of the first ST1/RT027 isolate in 1985 (*2, 3*). He and colleagues (2013) analyzed a dataset of ST1 genomes and described two epidemic lineages, both of which acquired the GyrA Thr82Ile mutation conferring quinolone resistance; the time of emergence for these two ST1 lineages was estimated to be in the early 1990s. The SNP accumulation rate within *C. difficile* must be considered when attempting to track isolates on an epidemiological time scale (*103*) and cases linked by recent transmission will likely be separated by very few SNPs (*26*). The genomic variation among northern Arizona isolates suggests that although some CDI cases from which these isolates were collected may potentially be the result of transmission within healthcare facilities, other cases are seemingly unrelated. We observed that closely related isolates, based on WGS analysis, were sampled in patients at multiple facilities and over multiple time points. In two instances, whole genome sequencing showed that the same genotype persisted in a patient rather than the patient being infected with a different genotype. Since patients with CDI are moved between facilities and similar *C. difficile* genotypes are detected at different facilities, infection prevention strategies must be implemented across the entire healthcare network to be most effective. Putatively unrelated cases may be the result of particular ST1 genotypes circulating in northern Arizona reservoirs and emerging as CDI cases due to antibiotic use or other factors. Importantly, monitoring transmission and persistence of these ST1 isolates with sub-genomic methods, such as ribotyping or MLST, would not have been possible due to a lack of resolution provided by these methods. Incorporating whole genome sequencing and comparative genomics into the healthcare network surveillance program enabled more effective monitoring, which can hopefully improve patient health in the future.

AMR varies within *C. difficile* (*36*) and AMR markers detected in genomes for ST1 isolates sampled in this study indicated that *in silico* predicted AMR profiles for these isolates are also variable. Resistance to fluoroquinolones within *C. difficile* ST1 has been the focus of many studies (*4, 9, 37–39*) and approximately 95% of the ST1 genomes compared in this study have the missense mutation in the *gyrA* gene (Figure 2, Table S9) linked to resistance to some fluoroquinolones. The use of quinolones to treat un-related infections may select for ST1, supporting the proliferation of clinical CDI caused by this sequence type. Interestingly, the GyrA Thr82Ile mutation and other mutations known to confer quinolone resistance were absent in five of the 27 ST1 isolates collected in this study. These five isolates account for four introductions of the ST1 lineage into the healthcare network in this region (clades 2-5 in Figure 2); one facility within the healthcare network implemented a program to restrict fluoroquinolone use in 2017, which may reduce selection for the GyrA Thr82Ile genotype. AMR screening of northern Arizona isolates with varying *in silico* predicted AMR profiles (Table S10) indicated that all isolates were resistant to ciprofloxacin; resistance to ciprofloxacin is common among *C. difficile* (*36*). Resistance to tetracycline was indicated for one northern Arizona isolate for which a *tetM* gene was identified in the genome. Use of tetracyclines has not been frequently associated with CDI (*104*), but a recent report has associated tetracycline resistance to RT078 (*105*). No strains were fully resistant to vancomycin, which is a recommended treatment option for CDI (*42*), despite the presence of CDSs associated with vancomycin resistance being detected in one tested genome. Continued screening of *C. difficile* antimicrobial resistance is important to monitor AMR variability and provide insight for updating treatment guidelines.

Although AMR plays an important role in *C. difficile* epidemiology, other factors, in addition to quinolone resistance, likely contribute to the prevalence of CDI attributed to ST1. For example, the GyrA Thr82Ile mutation is also highly conserved in ST63 (RT053) genomes, although this ST is not as successful as ST1 based on reported genotype frequencies from hospital isolation (*11, 13, 26, 27*). Additionally, *C. difficile* ST1 appears to be less prevalent than other STs in the natural environment (samples not associated with the human gut or hospital) based upon environmental survey studies (*30, 31, 106*). LS-BSR analysis indicated that genomic regions were highly conserved within ST1 genomes, as well as a number of regions that have been broadly deleted within ST1 genomes (Table S12). Reduction of the core genome can be associated with niche differentiation (*107, 108*). The observed genome region acquisition and loss across ST1 may be associated with its almost exclusive presence in the human/hospital environment. An analysis of other STs demonstrates that the gene gain/loss may be more prevalent in ST1 than other toxigenic clades. However, as most sampling efforts are focused on clinical settings, prevalence of ST1 in other environments is not fully understood. Genetic variability within ST1 may also contribute to the success of the lineage. For instance, resistance to clindamycin is not universal within ST1 (*4, 109*) but would provide an advantage to some ST1 isolates. Genomics alone cannot definitively identify why ST1 is such a successful lineage of *C. difficile*, but can identify targets for further investigation (e.g. CDSs identified in Table S12). Additionally, here we focus on bacterial genomics without consideration of factors such as variation in gene regulation, host immune response, or hospital practices such as antimicrobial use policies. For example, with the curtailed use of quinolones in some areas, ST42 (RT106) now appears to be the most dominant *C. difficile* sequence type in the United States (*110*).

*C. difficile* isolates have been differentiated using a variety of methods including MLST and PCR ribotyping. In this study, we used the MLST designation, as we can easily genotype sequenced genomes *in silico* in the absence of ribotyping data. However, due to recombination, phylogenies inferred from a concatenation of MLST markers have been shown to be incongruent with WGS phylogenies (*111*). Using whole genome sequence data for *C. difficile*, we demonstrate that multiple STs are paraphyletic on the core genome SNP phylogeny (Figure 1). This is important if common STs are expected to have a shared evolutionary history, yet the history is conflicting when using the entire genome. Ribotyping represents a legacy method that is still used to type isolates and infer relationships. However, there remains a disconnect between ribotyping and sequencing methods. In this manuscript, we present a workflow that can associate raw sequence data from ST1 genomes to a ribotype profile (RT027). The amplicons that were used for *in silico* ribotyping were identified from only three complete ST1 genomes; draft genomes are not useful for the identification of predicted amplicons due to the potential collapse of repeat regions (e.g. 16S rRNA genes) (*112*). As additional complete genomes are generated from diverse ribotypes, this method can be modified to associate ribotype information with whole genome sequence and MLST data, which is useful for bridging between different methodologies. Whole genome sequencing and analysis provides the highest resolution method for comparing *C. difficile* isolates and should be included in epidemiological studies if possible.

This study has provided a large-scale analysis into a single sequence type of *C. difficile* in order to place isolates from a local healthcare network into a global context and to understand the diversity within a successful, world-wide *C. difficile* sequence type. The analyses indicated multiple introductions of *C. difficile* ST1 into the healthcare network, provided insight into potential transmission of *C. difficile* within the healthcare network, and identified AMR variation among isolates. The genomic diversity within the ST1 genomes suggests that broadly applying labels to ST1 genomes, such as hypervirulent clone, may be inappropriate without an in-depth analysis of the gene presence/absence and SNP markers for antimicrobial resistance. Additional studies into how genomic variation affects toxin production and virulence are needed to assess the phenotypic diversity of ST1 as it relates to the observed genotypic diversity; these studies are currently ongoing and will help shape how we approach studies using sub-genomic information, such as ribotyping and MLST.

## 5. Author statements

### Ethical approval

Stool samples identified as containing *C. difficile* from Northern Arizona Healthcare facilities were collected under IRB No. 764034-NAH.

## Acknowledgements

Computational analyses were run on Northern Arizona University’s Monsoon computing cluster, funded by Arizona’s Technology and Research Initiative Fund.

## Supplementary Figures and Tables

(Available from github: https://github.com/chawillia/cdiff_data)

Figure S1. Phylogeny of ST1 genomes paired with a heatmap indicating BSR values of a subset of proteins that are differentially conserved throughout ST1. Red arrows indicate examples of clade-specific genes/proteins, and the gray arrow indicates an example of genes/proteins that are deleted in a specific clade.

Figure S2: Bioanalyzer gel image for PCR ribotyping of *C. difficile* isolates. Lanes identified with blue boxes include ST1 isolates. The ST1 isolates have similar PCR ribotyping profiles, which are comparable to *in silico* ribotyping results (see main text).

Table S1: External genomes analyzed in this study

Table S2: NASP matrix of SNPs for global collection of *C. difficile*

Table S3: NASP matrix of SNPs for *C. difficile* ST1 genomes

Table S4: SNPs for timing analysis

Table S5: Assessment of statistical fits for clock and demographic model combination

Table S6: Previously described genomic regions associated with virulence in *C. difficile* ST1

Table S7: Ribotype region information for three complete *C. difficile* ST1 genomes

Table S8: Results for screening genomes for antimicrobial resistance markers (CARD database) with LB-BSR

Table S9: Summary of GyrA and GyrB mutations in *C. difficile*

Table S10: Results for antimicrobial resistance screening with Etests

Table S11: The strict core genome identified with LS-BSR for *C. difficile* ST1 genomes

Table S12: CDSs identified as ‘gained’ and ‘lost’ within *C. difficile* ST1

Table S13: Results for screening genomes for sequences or markers associated with *C. difficile* virulence

Table S14. *In silico* ribotyping threshold summary

Table S15: *In silico* ribotyping results for all analyzed genomes

Table S16: Comparison of *in silico* and PCR ribotyping results

